# Close genetic relationships between a spousal pair with autism-affected children and high minor allele content in cases in autism-associated SNPs

**DOI:** 10.1101/073452

**Authors:** Zuobin Zhu, Xitong Lu, Dejian Yuan, Shi Huang

## Abstract

Parents of children affected with autism spectrum disorders (ASD) often have mild forms of autistic-like characteristics. Past studies have focused on searching for individual genetic risk loci of ASD. Here we studied the overall properties of the genomes of ASD trios by using previously published genome-wide data for common SNPs. The pairwise genetic distance (PGD) between a spousal pair with ASD-affected children was found smaller than that of a random pair selected among the spouses in the ASD trios, and spousal relatedness correlated with severe forms of ASD. Furthermore, for a set of 970 ASD associated SNPs, cases showed higher homozygous minor allele content than parents. These results indicate new genetic elements in the broad phenotypes of parents with ASD-affected offspring and in ASD pathogenesis.

**Highlights:**

- The overall properties of the genomes of ASD trios show ASD specific features.
- A spousal pair with ASD-affected children showed smaller genetic distance than that of a randomly selected pair among the spouses in the ASD trios.
- Minor allele contents in ASD-associated SNPs in ASD cases were greater than those in the parents of cases.

## Introduction

Autism spectrum disorders (ASD) is a common disease today. The prevalence of ASD is increasing and reached 14.6 per 1,000 (one in 68) children aged 8 years in the United States at 2012[1]. About four times as many males as females are autism [2; 3]. Twin and family studies show that siblings of children with ASD are at a significant higher risk for autism than the general population[4; 5; 6]. Strict (STR) and spectrum (SPC) definition of ASD differ mainly in social deficits[7]. Parents of ASD children are in general of high social economic status (SES) with similar background in science and engineering fields but often have mild forms of autistic-like characteristics or the ‘broad phenotype’ of autism, such as social and communicative difficulties[8; 9; 10; 11]. High SES and educational attainment are strongly correlated, both of which are also correlated with general intelligence[12; 13]. There is also a genetic component to educational attainment[14]. ASD children also show wide distribution in general intelligence with high functioning individuals performing better than the general population or even their high SES parents in certain tasks[15; 16; 17].

ASD remains poorly understood but may have a strong genetic component with a heritability of 40–80%[18; 19; 20; 21]. ASD are genetically highly heterogeneous, with no single gene accounting for more than 1% of cases[22]. Recent work has shown a substantial contribution of de novo variations[23; 24; 25; 26]. However, genome-wide association studies have revealed few replicable common polymorphisms associated with ASD[27; 28; 29; 30].

Theories of ASD are numerous. According to the hyper-systemizing theory[31; 32], people with ASD have an unusually strong drive to systemize. A comprehensive hypothesis of ASD, taking intelligence and nearly all aspects of ASD into account, has emphasized the role of an optimum level of a suppressive force of innate traits[12].

It has been reported that similar phenotypes and genetic parameters influence preferential mating[33; 34; 35]. Consanguineous marriages appear to increase the prevalence of ASDs[36; 37; 38]. The prevalence of autism might increase by 1.5-fold after 1 generation of assortative mating (≥2.4-fold in the long term) depending on several assumptions[39]. Common genetic variants are individually of little effect but may be a major source of risk for autism[40]. Preferential mating may bring about additive genetic influences in concentrating inherited ASD susceptibility[41]. These observations suggest a potential role for combination of common variants in ASD.

Consistent with the notion of a collective and additive effects of common variants, recent studies indicate a role for genome wide minor allele content (MAC) of an individual in a variety of complex traits and diseases[42; 43; 44; 45]. The more the number of minor alleles of common SNPs in an individual (i.e., the higher the MAC values), the higher the risk in general for many complex diseases such as lung cancer and Parkinson’s disease[42; 43; 44; 45]. Such findings indicate an optimum level of genetic variations that an individual can tolerate. Too much lower or higher than the optimum may result in lower fitness and complex diseases[46; 47]. In this study, we investigated whether spousal pairs with ASD-affected children are more genetically alike and whether changes in MAC values may be linked with ASD.

## Results

### Genetic similarities between a spousal pair with ASD-affected children

To determine whether the spousal pairs with ASD-affected children may be genetically more related to each other, we calculated the pairwise genetic distance (PGD) between a spousal pair with ASD-affected children relative to the PGD between a randomly chosen pair of male and female within the parents population in the Autism Genome Project (AGP) cohort[48; 49] and in the Miami dataset[50]. The AGP consists of two cohorts, Stage 1 and Stage 2. As controls for the spousal pairs in the ADS trios, we used the control cohort from the Australian twin-family study of alcohol use disorder (OZALC study)[51] and the European family in 1000 genomes project. As general population controls for the AGP ASD trios. We made use of the published SNP genotyping data from these autism and controls studies, and only used shared autosome SNPs for comparative analyses involving different datasets. We performed PCA analysis of control cohorts in combination with the autism cohorts (Supplementary Fig. S1). To minimize population heterogeneity, we only selected individuals that form a relatively tight cluster as shown in Supplementary Fig. S1.

We used two methods to calculate pairwise genetic relationships. The first method measures PGD and made use of a previously described custom script that does not take into account the frequency of the SNPs in the population: all the SNPs contributed equally to genetic distance[45]. For homozygous (Hom) vs Hom mismatch, a difference of 1 was scored. For Hom vs Het or for Het vs Het, a difference of 0.5 was scored. The score of 0.5 for Het vs Het match can be intuitively understood as described in detail in the Methods section. Common software for calculating genetic distance such as PLINK and PEAS typically score Het vs Het as 0 distance, which fail to take into account of haplotype differences and are hence not realistic[52; 53].

To generate random pair distance, we calculated the distance between a father in a parental pair and each of the female individuals in the parents cohort. We then compared the parental pair distance with the middle ranked distance among the randomly paired distances. If the parental pair is indeed closer than a random pair, its distance should be smaller rather than larger or similar to middle ranked random pair distance. We did this for all parental pairs and performed pairwise T test to determine whether the parental pair distance is significantly smaller than the middle ranked random pair. We found that the PGD of a spousal pair was significantly smaller than that of a random pair in both Stage 1 (*P* = 1.27E-9) and Stage 2 (*P* = 8.71E-18) cohorts of the AGP. Similar trend was found in the Miami dataset (*P* = 0.08) (Fig. 1a). However, the same type of analysis did not reveal a difference between spousal pairs and random pairs in the general population OZLAC and CEU (Fig. 1a).

**Fig 1.**
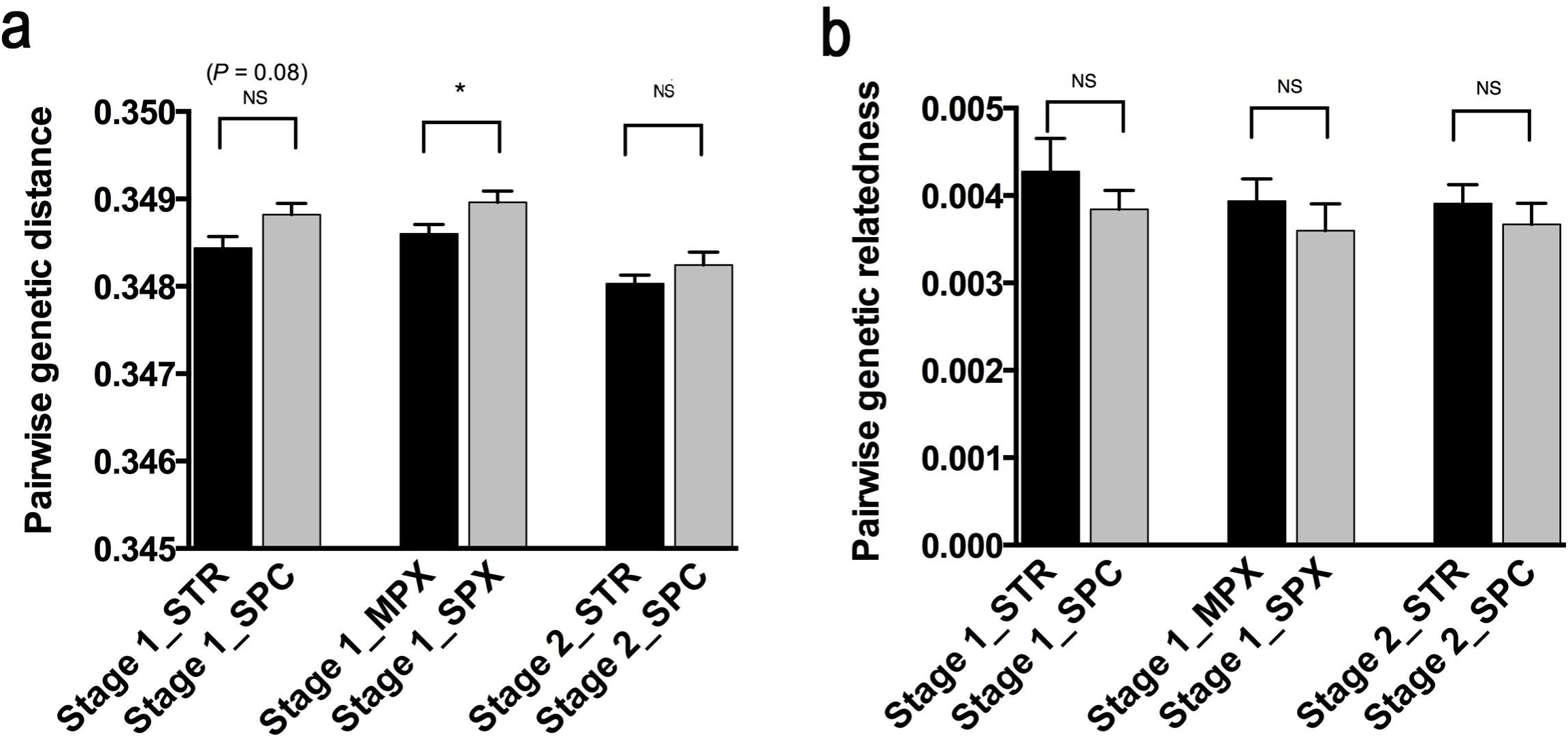
**Pairwise genetic distance (PGD) between spousal pairs and between randomly selected pairs.** PGD of spousal pairs vs random pairs using either Stage 1 or Stage 2 dataset of AGP and the Miami dataset. Control spouses were from the Australian twin-family study of alcohol addiction study (OZALC) and the CEU data from 1000 genomes. (a) PGD calculated by the custom script. (b) Genetic relatedness calculated by the GCTA method. *** *P* <0.001, NS *P* > 0.05, paired T test. Data are means ± S.E.M.

We repeated the experiments by employing an alternative measure of pairwise genetic relatedness, the equation 3 of genome wide complex traits analysis(GCTA) that takes into account the frequency of alleles in the population: sharing a rare allele is given a higher weight for genetic relatedness compared to sharing a common allele[54]. This method generates relatedness value rather than distance value with higher value meaning closer genetic distance. Using this relatedness index, we again obtained the same results showing parental pairs having higher relatedness values than random pairs (Stage 1 dataset, *P* = 0.0009; Stage 2 dataset, *P* =0.0007; Miami dataset, *P* =0.06) (Fig 1b). Here, different datasets from independent studies gave different relatedness values, which may be related to sample size variations since the GCTA method makes use of allele frequency the reliability or accuracy of which is dependent on sample size. Small population size would gave inaccurate allele frequency estimation and may make more alleles appear as rare, which would lead to overestimation of genetic relatedness by the GCTA method. The results in Fig. 1b did show a trend of more relatedness in datasets with small population size. Likewise, sample size would also affect estimation of average PGD as shown in Fig. 1a with small sample size being less accurate. So, average PGD between independent datasets may not be identical.

To study if the relatedness of parents correlates with disease status, which would also serve to verify that it was not a result of trivial bias during genotyping experiments such as batch effects, we compared the PGD of a spousal pair with strictly defined (STR) ASD-affected children with that of a spousal pair with spectrum defined (SPC) ASD-affected children, expecting SPC cases to be relatively more control like. We also compared the PGD of a spousal pair with multiple incidence (multiplex, MPX) of ASD-affected children with that of a spousal pair with single incidence (simplex, SPX) of ASD-affected children, expecting the SPX cases to be relatively more like the controls. For Stage 1 data of AGP, by using the first method, we found the PGD of a spousal pair with STR (*P* = 0.08) or MPX (*P* = 0.04) affected children to be weakly smaller than that with SPC or SPX affected children (Fig. 2a). For Stage 2 data, we did not find significant difference regarding STR and SPC comparisons and there were not enough number of cases to perform MPX versus SPX comparisons (Fig. 2a). The Miami dataset was not suitable for this type of analysis as it had few numbers of relevant cases. By using the second method, we did find the same trend although insignificant (Fig 2b), indicating that allele frequency based distance method may not be sensitive enough in cases where the cohort was relatively small as in the case here with allele frequency less accurate or realistic. Overall, these results indicate close genetic relationship between a spousal pair with ASD children, and a specific association of this relationship with ASD severity.

### Minor allele contents in ASD-associated SNPs in ASD cases

It has been reported that Parkinson’s disease (PD) population has higher MAC than control population[44] and that mice with more heterozygosity (Het) show more sociability than mice with less[55]. Here we tested these genome wide characteristics in the ASD trios and control populations. We calculated the amount of homozygous SNPs, homozygous minor alleles content (homoMAC) and MAC in the ASD trios of AGP and the Miami dataset and the control populations. However, we found no difference in these values between the parents and children in the datasets of either cases or controls (Fig. 3).

To test whether a set of risk SNPs may reveal a difference between parents and their case children, we next studied the amount of homozygous SNPs, homoMAC and MAC using the risk SNPs located in candidate ASD-associated genes as reported by the literature or the SFARI database[56]. We found 1114 SNPs located in the ASD-associated genes. Using the pairwise linkage disequilibrium (LD) option of PLINK, we calculated the LD of each neighboring SNPs, and found 124 SNPs in LD with their neighboring SNPs with R2 > 0.5. Most of these LD groups involved on 2 SNPs and most of 1114 SNPs examined here were not in LD. We randomly selected one among SNPs in LD in order to use only 1 SNP to represent each haplotype. Thereby, we obtained a final set of 970 SNPs as ASD-associated SNPs (Supplementary Table S1). We found no difference in the amount of homozygous SNPs between parents and children (Fig 4a). However, homoMAC values in these SNPs were higher in ASD cases relative to that of their parents (significant in Stage 1 data, *P* =0.006; and Stage 2 data, *P* = 0.003; insignificant in the Miami data) (Fig 4b). Also, ASD case children in the Stage 1 (*P* = 0.0003) and 2 (*P* = 0.002) and the Miami dataset (*P* = 0.006) had higher MAC than their parents (Fig 4c). We did not find a relationship between homoMAC or MAC and disease severity. Thus, MAC in a small set of risk SNPs located in candidate ASD genes may play a role in ASD disease onset but not in progression.

## Discussion

In this study, we found closer genetic similarities between a spousal pair with ASD children than that between randomly paired males and females. Such relatedness of parents correlated with disease status. Furthermore, relative to parents, cases were linked with higher homozygous MAC in ASD-associated SNPs. These results were robust and repeated using two different distance scoring methods and 2 or sometimes 3 ASD datasets.

A caveat with our study involving different datasets from different sources is the potential bias introduced by non-uniform genotyping analyses. We have controlled for this by using only shared SNPs among different datasets and only SNPs with normal frequency distribution consistent with the Hardy Weinberg equation. We used middle ranked distance among all possible random pairs rather than average distance. This should minimize the impact of population outliers on comparing parental pair distance and the distance between random pairs. That ASD spousal pairs are more related is unlikely a result of trivial genotyping batch effects, since spousal relatedness also correlated with disease status. The key findings of this study did not involve any control dataset and was based on the same set of uniformly genotyped data, which include closer ASD spousal pair distance relative to random pairs, correlation of spousal relatedness with disease status, and higher homozygous MAC in ASD-associated SNPs in the ASD-affected children relative to their parents.

A previous study has found a closer genetic similarity between a spousal pair in the general population relative to between randomly paired males and females[34]. However, we did not repeat this observation in our control population. It is likely that even if spousal pairs in the general population are more related than random, it may not be as robust as the ASD spousal pairs. Consistently, ASDs have been found to be more common in consanguinity marriages[36; 37; 57]. The AGP samples were mostly from Europe and some from Canada (about 85% of AGP samples were European ancestry with the remainder other ancestries) while the controls we used were mostly European Americans who may be expected in general to be more genetically diverse than Europeans. We have corrected for this by using PCA analysis to exclude outliers. Regardless, however, these demographic factors may not affect the key findings as listed above that did not involve controls and used only AGP samples.

Our previous studies show increased genome wide MAC values in patients with Parkinson’s disease and several other types of complex disease[42; 43; 44; 45]. However, the genome wide MAC values of ASD-affected children were not found to be significantly different from their parents. Nonetheless, they did show increased homoMAC than their parents in ASD-associated SNPs, consistent with a more deleterious role for minor alleles in ASD pathogenesis. The difference in average homoMAC between cases and parents seems small and yet significant. This is consistent with previous findings on PD and other complex traits and diseases[44; 45; 58]. The optimum MAC level may be very finely balanced and slightly abnormal traits such as ASD or PD may result with just a slight excess in MAC.

The finding here strengthens the concept of a genome wide optimum level of genetic diversity[46; 47]. There may exist an optimum level of genetic variations for a set of genes responsible for a particular trait. An increase over such a trait-specific optimum threshold may not necessarily bring about significant change in overall genome wide genetic diversity or MAC in an individual, which would depend on how many loci are involved in controlling a trait, but could nonetheless be sufficient to result in abnormality in the trait and hence disease. Overall, our results here suggest new genetic elements for ASD and the broad phenotypes of parents with ASD-affected children.

## Materials and methods

### Cohort description

Two GWAS family datasets of ASD, Stage 1 and Stage 2, from the Autism Genome Project (phs000267.v4.p2) and one GWAS family dataset from the Miami study (phs000436.v1.p1) were downloaded from database of Genotypes and Phenotypes (dbGaP)[48; 49; 50]. Controls derived from convenience samples. All the controls used in this study were more than 18 years old healthy adults. Similar to previous studies that used convenience controls[40], we reasoned that ASD is sufficiently rare (approximately 1.46%) that screened and unscreened controls would yield similar results[1]. As controls for the spousal pairs in the ADS trios, we downloaded from dbGaP the data on the spouses in the Australian twin-family study of alcohol addiction study (OZALC) (phs000181.v1.p1)[51]. As general population controls for the AGP ASD trios, we downloaded from dbGaP a control populations based on overlapping SNP genotypes and similar demographic profiles. All analyses were done with autosomal SNPs which ~300K were shared with all the datasets used here. Genotype distributions for each SNP were consistent with Hardy-Weinberg equilibrium (P > 0.01).

We only used shared SNPs for comparative studies between ASD and controls. The OZALC samples were genotyped on Illumina HumanCNV370v1 containing 370K SNPs, of which ~300K were shared with the AGP data and Miami data.

### Population Stratification

The AGP Stage 1 dataset comprised 1471 families, of which 1141 were previously identified to be of European ancestry[48; 49]. AGP Stage 2 dataset had 1301 families, of which 1108 were previously identified to be of European ancestry[48; 49]. The Miami dataset comprised of 377 Caucasian families[50]. Principal component analysis (PCA) using the GCTA tool was used to estimate the genetic relatedness[54]. The shared ~300K SNPs that have passed quality control filters were used here. We excluded the outliers by principle component 1-3 (Supplementary Fig. S1). Duplicated individuals were excluded from the analysis. After outlier exclusion, there were 320 spouses in AGP stage 1 trios, 283 spouses in AGP stage 2 trios, 133 spouses in Miami ASD trios, 125 spouses in OZALC data and 22 spouses in CEU data (Supplementary Fig S1).

### Comparison of parental pair distance with a random pair distance

To generate random pair distance for comparison with a parental pair, we calculated the distance between a father in a parental pair and each of the female individuals in the parents cohort. We then compared the parental pair distance with the middle ranked distance among the randomly paired distances. We did this for all parental pairs and performed pairwise T test to determine whether the parental pair distance is significantly smaller than the middle ranked random pair. The middle ranked distance among the random pairs, relative to average distance of random pairs, should be less sensitive to influence by very large or very small distances from certain random pairs that may be due to paring of demographically very distant or very close individuals that may represent a small fraction of our selected cohort despite our best effort to select homogeneous populations. For example, 20 outliers in an otherwise homogeneous cohort of 300 may significantly raise the average random pair distance but would only marginally affect the middle ranked random pair distance.

### Selection of SNPs within ASD-associated genes

The ASD associated genes were obtained from the published papers^49^. We first selected the ASD-associated SNPs as those that are located within the ASD associated genes, or within 1,000 base pairs upstream or downstream and the UTR regions of these genes. Using the pairwise linkage disequilibrium (LD) option of PLINK, we calculated the LD of each neighboring SNPs (the LD window was 1 million base pairs on the same chromosome). If there were two or more SNPs in LD with R² > 0.5, we randomly selected one among these SNPs.

### Statistical analysis

The population used for calculating the pairwise genetic distance (PGD) were homogeneous groups with outliers excluded by “GCTA” (genome-wide complex trait analysis). PGD were scored using a software as described in previous studies^43,44^. Every non-repetitive pair within a population was scored to produce the average PGD. The PGD software measures genetic distance between two individuals by the number of mismatched SNPs. For homozygous (Hom) vs Hom mismatch such as CC vs TT, a difference of 1 was scored. For Hom vs Het such as CC vs CT, a difference of 0.5 was scored. For Het vs Het such as CT vs CT, a difference of 0.5 was scored, which is based on the following reasoning. When there is AB v AB match, there are two situations depending on the haplotypes. First, if haplotypes are matched, the two hets would be identical (AB matched with AB) and it would take 0 mutation to convert AB to AB. Second, if haplotypes are not matched, AB would be matched with BA. It would take 2 mutations to convert AB into BA. Since only 50% of het vs het matches would be AB vs BA, so the overall number of mutations required to make AB and BA equal to AB in terms of haplotype matches is 0.5 × 2 = 1, which is 50% lower than that required for changing AA to BB. Since we score AA v BB as a difference of 1, the score for AB v AB is naturally 0.5. We verified this approach by comparing the PGD in X chromosome for CEU females vs CEU males using HapMap SNP data and found them to be similar as expected. In contrast, a software based on IBS (identical by status) such as PEAS that score A/B vs A/B as 0 showed the males to have much greater PGD in X than females[53]. For the missing genotypes N/N, N/N vs Hom was scored as 0 and N/N vs Het as 0.5. All the PGD (or the ratio of homozygous genotype) were expressed as total number of the distance (or homozygous SNPs) divided by the total number of SNPs that actual used except the N/N.

The MAF of each SNP was calculated by PLINK and SNP Tools for Microsoft Excel[52; 59]. From MAF data of controls we obtained the MA set, which excluded non-informative SNPs with MAF = 0 in both cases and controls or with MAF = 0.5 in controls.

Minor allele content (MAC) means the ratio of the number of minor alleles divided by the total number of SNPs scanned.

## Acknowledgements

This work was supported by the National Natural Science Foundation of China grant 81171880 and the National Basic Research Program of China grant 2011CB51001 (S.H.).

The datasets used in this manuscript were obtained from dbGaP at http://www.ncbi.nlm.nih.gov/sites/entrez?Db=gap. The dbGaP accession numbers include phs000267.v4.p2, phs000436.v1.p1and phs000181.v1.p1. We thank the Contributing Investigator(phs000267.v4.p2, Stephen W. Scherer, Bernie Devlin,etc; phs000436.v1.p1, Margaret A. Pericak-Vance; phs000181.v1.p1, Andrew C. Heath, Richard D. Todd, Pamela A. Madden, Alexandre Todorov etc. The Autism Genome Project (AGP) Consortium were supported by the National Institutes of Health, Bethesda, MD, USA, (phs000267.v1.p1). (HD055751,HD055782, HD055784, HD35465, MH52708, MH55284, MH57881, MH061009, MH06359, MH066673, MH080647, MH081754, MH66766, NS026630, NS042165 andNS049261). The Miami study were supported by the National Institutes of Health, Bethesda, MD, USA(AUT R01 R01MH080647 and AUT PPG P01NS026630) and Hussman Foundation(Gift). The Australian twin-family study of alcohol use disorder (OZALC study) were supported by HHSN268200782096C. NIH contract “High throughput genotyping for studying the genetic contributions to human disease”. National Institutes of Health, Bethesda, MD, USA(phs000181.v1.p1). The content is solely the responsibility of the authors and does not necessarily represent the official views of the funding agencies.

## Figure Legends

**Fig. 2. Genetic distance of spousal pairs with STR, SPC, MPX or SPX-affected children.** STR: strictly defined ASD, SPC: spectrum defined ASD, MPX: multiplex families, SPX: simplex families. (a) Pairwise genetic distance (PGD) calculated by the custom script. (b) Genetic relatedness calculated by the GCTA method. \**P* <0.05, NS *P* > 0.05, Z test. Data are means ± S.E.M.

**Fig. 3. Genomic characteristics in ASD cases and their parents.** (a) Shown are proportions of homozygous SNPs of parents or ASD-affected children in either Stage 1 or Stage 2 dataset of AGP, the Miami dataset, and the two control datasets. (b) Shown are proportions of homozygous minor allele genotype of parents or children. (c) Shown are minor allele content (MAC) of parents or children. NS *P* > 0.05., Z test. Data are means ± S.E.M.

**Fig. 4. The proportion of homozygous risk SNPs, homozygous minor allele contents and MAC in ASD trios.** (a) Shown are proportions of homozygous SNPs of parents or ASD-affected children in either Stage 1 or Stage 2 dataset of AGP, the Miami dataset, and the two control datasets. (b) Shown are proportions of homo MAC of parents or children. (c) Shown are MAC of parents or children. ** *P* <0.01, ** *P* <0.01, NS P > 0.05, Z test. Data are means ± S.E.M.

